# Parthenogenomics: insights on mutation rates and nucleotide diversity in parthenogenetic *Panagrolaimus* nematodes

**DOI:** 10.1101/2022.09.15.508068

**Authors:** Laura I. Villegas, Luca Ferretti, Thomas Wiehe, Ann-Marie Waldvogel, Philipp H. Schiffer

## Abstract

Asexual reproduction is assumed to lead to the accumulation of deleterious mutations, and reduced heterozygosity due to the absence of recombination. Panagrolaimid nematode species display different modes of reproduction. Sexual reproduction with distinct males and females, asexual reproduction through parthenogenesis in the genus *Panagrolaimus*, and hermaphroditism in *Propanagrolaimus*. Here, we compared genomic features of free-living nematode populations in populations and species isolated from geographically distant regions to study diversity, and genome-wide differentiation under different modes of reproduction. We firstly estimated genome-wide spontaneous mutation rates in a triploid parthenogenetic *Panagrolaimus* population, and a diploid hermaphroditic *Propanagrolaimus* species via long-term mutation-accumulation-lines. Secondly, we calculated population genetic parameters including nucleotide diversity, and fixation index (*F*_*ST*_) between populations of asexually and sexually reproducing nematodes. Thirdly, we used phylogenetic network methods on sexually and asexually reproducing *Panagrolaimus* populations to understand evolutionary relationships between them. The estimated mutation rate was slightly lower for the asexual population, as expected for taxa with this reproductive mode. Natural polyploid asexual populations revealed higher nucleotide diversity. Despite their common ancestor, a gene network revealed a high level of genetic differentiation among asexual populations. The elevated heterozygosity found in the triploid parthenogens could be explained by the third genome copy. Given their tendentially lower mutation rates it can be hypothesized that this is part of the mechanism to evade Muller’s ratchet. Our findings in parthenogenetic triploid nematode populations seem to challenge common expectations of evolution under asexuality.

## 1 Introduction

Parthenogenetic organisms reproduce asexually, without paying the cost of sex, i.e. production of males, courtship or mate finding. Parthenogenetic reproduction should result in an evolutionary advantage (Otto & Lenormand, 2002), as all-female species are in theory capable of generating more offspring from the same resources, when compated to sexual sister species. However, sexual reproduction is the predominant reproductive mode among Metazoans, leading to a paradox that was coined as the “Queen of Evolutionary questions” by Graham Bell (Bell, 1982). To explain the apparent “lack” of parthenogenetic taxa, evolutionary theory posits that the absence of recombination inhibits the efficient purging of deleterious mutations which accordingly accumulate over time. This effect, called Muller’s ratchet (or mutational meltdown e.g. for RNA viruses) (Gabriel, Lynch, & Burger, 1993) (Felsenstein, 1974) counteracts the long-term persistence of obligate asexual lineages lacking recombination. Hence, mutation rates, the rate at which mutations arise *de novo* in a genome, are thought to vary between organisms with different reproductive modes, being lower in asexually reproducing organisms when compared to sexual relatives (Sloan & Panjeti, 2010).

Additionally in parthenogens, due to the lack of outcrossing, alleles can’t be recombined and the linkage between sites in finite populations would reduce the overall effectiveness of selection (Hill & Robertson, 1966). The Red Queen Hypothesis (RQH), originally focused on coevolution in a host-parasite context, has been extended to propose that sexually reproducing organisms have an evolutionary advantage in habitats with many biotic interactions, while asexually reproducing taxa are expected to be more frequent in habitats with challenging environmental conditions and lowered biotic pressure (e.g. higher altitudes) (Hartmann et al., 2017; Van Valen, 1973). Empirical studies paint a different, more complex picture of the evolutionary persistene of parthenognes: Bdelloid rotifers, oribatid mites and Darwinulid ostracods are examples of the so called ‘evolutionary scandals’, asexual taxa apparently persisting on geological time-scales. (Smith, 1978). Despite the expected reduced genetic diversity in parthenogenetic taxa, a study on the triploid asexual crayfish *Procambarus virginalis*, has shown that this species is highly heterozygous even when the expected outcome of its reproductive mechanism would be homozygosity (Schwarz, 2017). It might thus be possible for natural populations of asexually reproducing organisms to genetically diverge due to local adaptation, genetic drift, geographic isolation, and the generation of distinct mutations in different populations. Currently, there is still a lack of data on such asexually reprodcuing organisms and populations, in part due to potential study systems not lending themselves to easy cultivation in the laboratory for molecular evolutionary experiments.

Nematoda, one of the most species-rich groups within Metazoa, is a highly diverse phylum in terms of ecological niches inhabited, size ranges of different groups and reproductive strategies (Blaxter, 2011). The family Panagrolaimidae, the focus system of this study, exhibits various reproductive modes: gonochorism (sexual), parthenogenesis (asexual), and hermaphroditism (sexual, but selfing)(Kiontke et al., 2004). This system, with closely related taxa, can provide information on how genomic features may differ in relation to reproductive modes.

Panagrolaimidae nematodes have been used as a system to address diverse biological questions. Several panagrolaimids from distant regions (e.g. Russian Permafrost, Antarctica and Germany) are cryptobionts, displaying a suspended metabolic state (Lewis et al., 2009)(Shatilovich et al., 2023), providing a system to study and develop methodologies for long term storage of cells and tissues (Shatilovich et al., 2023). In addition, inactivation of gene expression in nematodes of the genus *Panagrolaimus* is also possible. RNA interference (RNAi) that could for functional genomic analysis has been succesful in panagrolaimids (Shannon, Tyson, Dix, Boyd, & Burnell, 2008). Recent studies have also performed gene editing in the genus mediated by CRISPR/Cas9 (Hellekes et al., 2023). Their varied reproductive modes, have allowed for the study of the evolution of parthenogenesis (Schiffer et al., 2019) (Shatilovich et al., 2023). Previous studies had suggested a common single origin of parthenogeneiss in the genus (Schiffer et al., 2019), however the newly described species *Panagrolaimus kolymaensis* has been proposed to have a second independent evolution of this reproduc. They are widely distributed across the globe, including extreme environments.

In this study, we made use of the *Panagrolaimus* system to to understand how mutation rates vary in free-living closely related species with different reproduction modes (sexual - selfing and asexual) and their genomic traits, such as nucleotide diversity and population divergence under pure inbreeding. We furthermore extended the study of these genomic traits to natural populations (here defined as a as a group of individuals from the same geographical origin belonging to the genera of interest) of diploid sexual and triploid parthenogenetic nematodes of the genus *Panagrolaimus*. We tested whether the results obtained in bottle-necked laboratory populations held true for natural Panagrolaimus populations from different geographical locations, and if these patterns of genome evolution in parthenogens are consistent with theoretical expectations or challenge them.

Figure 1. : organism picture.

**Figure 1:**
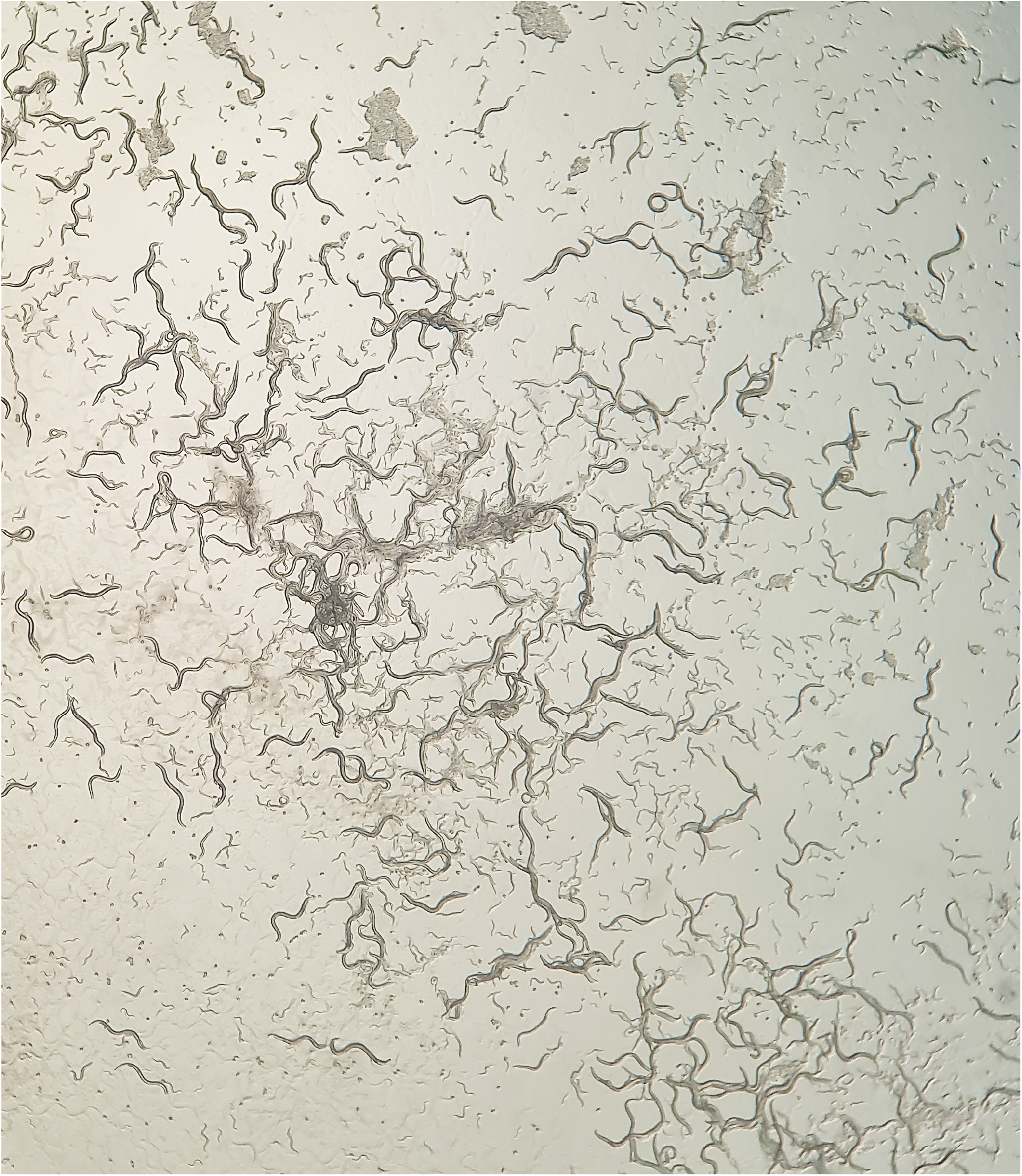
Panagrolaimidae is a family of nematodes from clade IV. Different reproductive modes can be found within the family: gonochorism, hermaphroditism and parthenogenesis.

## 2 Materials and Methods

### 2.1 Sampling, sequencing and data pre-processing

This study includes natural *Panagrolaimus* populations from different geographic locations that were previously described in McGill et al. 2015 (McGill et al., 2015) (supplementary table 1)(Figure 2). Asexual nematode populations used here are triploid (allopolyloids) and sexual populations are diploid selfing hermaphrodites, in which no males have been observed so far, reproductive modes of the nematodes used in this study have been previously defined in (Lewis et al., 2009) by single larval stage nematode propagation and determination of sperm presence using DIC/epifluorescence microscopy techniques. Populations were kept as laboratory cultures (at 15°C on low nutrient agar plates inoculated with OP50 *(E. coli)* from which DNA was extracted from several plates. Adult nematodes, as well as larvae and eggs were washed off from the plates and cleaned in 3 washing steps. After three rounds of freeze-thaw cycles on lysis buffer, genomic DNA was extracted following a salting-out genomic DNA protocol or using Qiagen’s genomic tip (cat.no. 10223). Whole genome sequencing of pooled specimens was performed on Illumina HiSeq2000 and NovaSeq platforms, sequencing data was deposited in SRA and is available under the Bioproject PRJNA374706 ([Dataset]Schiffer, 2017). After standard quality filtering and trimming of raw reads using fastp (versions 0.23.0 and 0.20.1) (Chen, Zhou, Chen, & Gu, 2018), paired-end reads were mapped to the respectively closest related reference assembly (available for populations PS1159, JU765, and ES5 (*P. sp*. ‘bornheim’),(supplementary table 2) using bwa-mem2 (version 2.2.1) (Vasimuddin, Misra, Li, & Aluru, 2019). For populations where reads were too short, mapping was done using NextGenMap (version 0.5.5) (Sedlazeck, Rescheneder, & von Haeseler, 2013). For populations where the insert size was smaller than double the read length, pear (version 0.9.8) (Zhang, Kobert, Flouri, & Stamatakis, 2014) was used before mapping with bwa-mem2. The alignments were filtered to remove duplicates using PICARD tools (MarkDuplicatesWithMateCigar)(version 2.26.8) (Institute, 2018), and lowquality reads (*<*30) were removed using samtools view (version 1.13) (Danecek et al., 2021). Asexual population PS1159 and sexual population JU765 were used in the mutation accumulation lines experiment for estimation of mutation rates. All other sexual and asexual populations were used in the population analysis and phylogenetic network. A list of the commands implemented in the following segments of Materials and Methods is provided here: https://github.com/lauraivillegasr/parthenogenomics.

**Figure 2:**
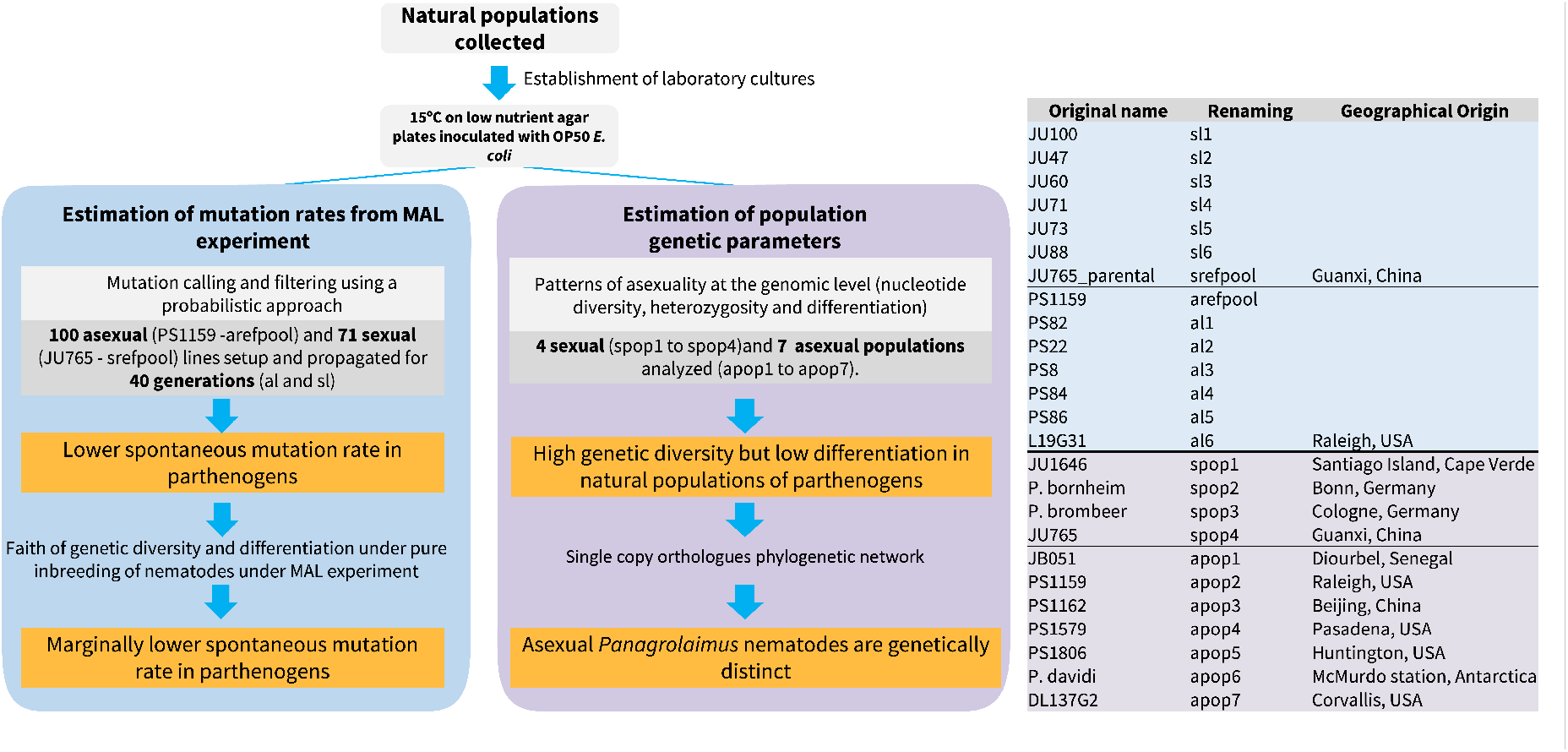
Workflow followed for this study. Populations used for the MAL experiment are highlighted in blue, populations used for estimation of population genetic parameters are highlighted in light purple. Main findigns of each section are highlighted in orange boxes.

### 2.2 Estimation of mutation rates from a MAL experiment

For the estimation of mutation rates, an asexual population *Panagrolaimus* sp. PS1159 and a sexual selfing population *Propanagrolaimus* sp. JU765 were subjected to a mutation accumulation lines (MALs) experiment for 30 to 52 generations of continuous inbreeding. Whole genome sequencing data was then generated from the starting point of the experiment and the end point of each MAL (Figure 3). DNA extraction was performed as described above. To allow for sufficient DNA content, several libraries of the respective population at the start of the MAL experiment were pooled into one, hereafter referred to as the *RefPool*. To ensure all quality scores were in Sanger encoding seqret (Madeira et al., 2019) and fastQC (version 0.11.9) (Andrews, 2010) were used. Mutations were called using a probabilistic approach (accuMUlate version 0.2.1)(Winter et al., 2018), following authors recommendations. Alignments from MAL were merged using samtools merge along with the RefPool. The analysis was performed only on a reduced set of positions commonly covered between all the lines and parental state per reproductive mode, respectively. Putative mutations were filtered by coverage ranges (332-575), number of reads supporting mutations (¿0.05), absence of mutant allele in other samples and absence of mutation in RefPool (==0), apparent mutation being caused by mismapped reads (Anderson-Darling test statistic) (¡=1.96), read pair successfully mapped to the reference genome (Fisher’s exact test)(specific commandsused for filtering can be found under: https://github.com/lauraivillegasr/parthenogenomics). Resulting candidates were manually curated using the Integrative Genomics Viewer (IGV version 2.16.1) (Thorvaldsdottir, Robinson, & Mesirov, 2013). The number of callable sites was estimated for each MAL as the number of positions along the assembly within the depth coverage of 10 and 50 (supplementary table 4). The mutation rates were obtained by dividing the number curated *de novo* mutations by the total number of callable sites using

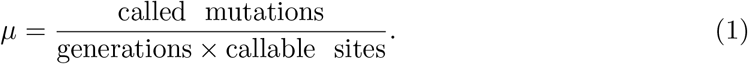

High-credibility intervals (HCI) for the estimated mutation rates were obtained using the Bayesian First Aid R package (version 0.2) (Bååth, 2014). Mutation rates estimated for each of the MAL, as well as the average mutation rate for both reproductive modes along with the average number of callable site per population, were provided as input for this.

**Figure 3:**
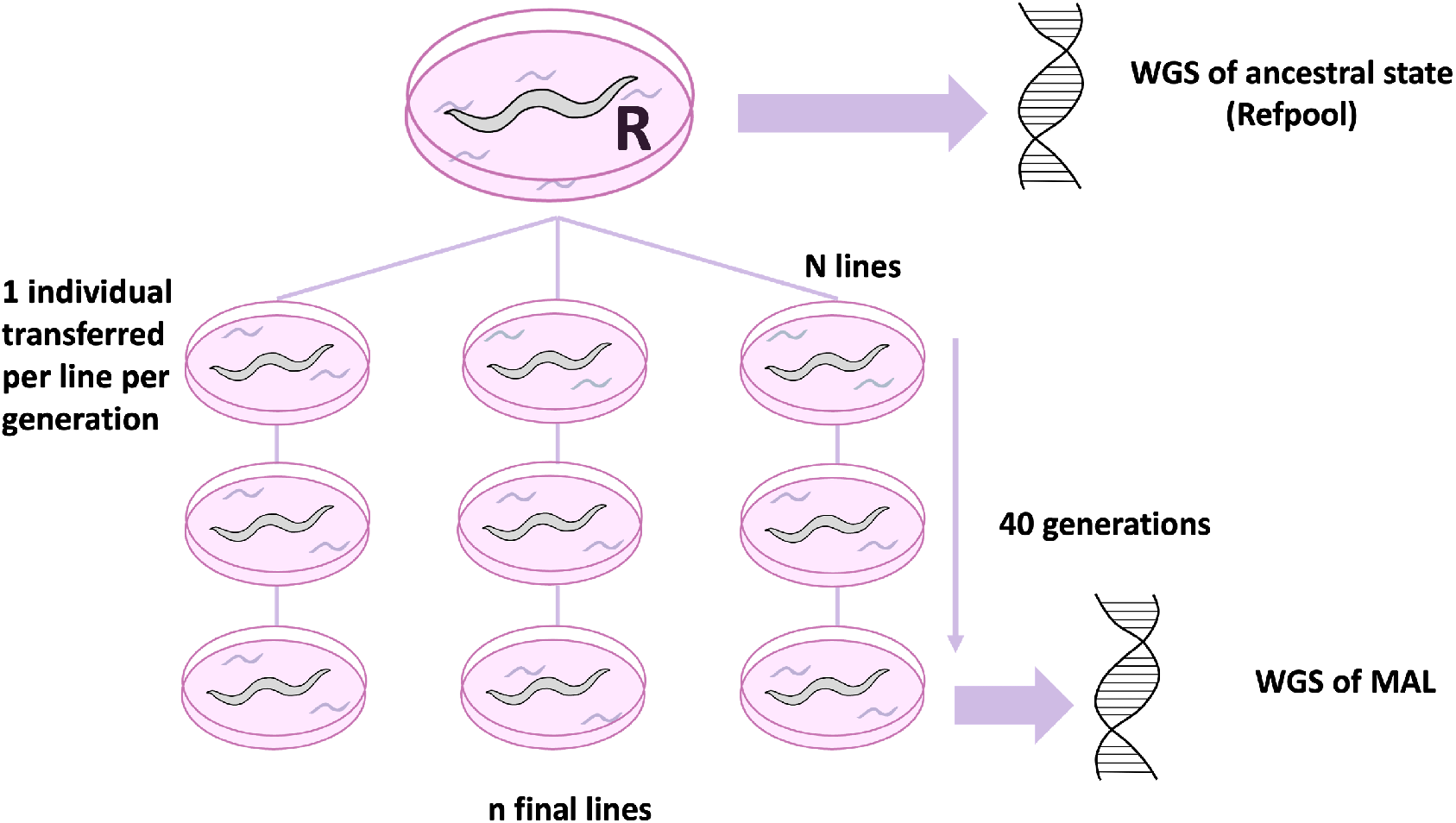
Experimental setup of mutation accumulation. A total of 100 asexual lines (PS1159) and 71 sexual lines (JU765) were established at the beginning of the experiment; the total of final lines was 15 for PS1159 and 30 for JU765.

### 2.3 Analysis of populations

#### 2.3.1 Population genetic parameters

Population genetic parameters *θ*_*w*_, nucleotide diversity (*π*), effective population size *N*_*e*_, and fixation index *F*_*ST*_ were estimated in two different settings (A) intrinsic estimation of *F*_*ST*_ in MA lines between each other and correspondent parental state (inbreeding escenario) and (B) between natural populations from different geographic origins (sexual populations were compared against asexual populations) . Both approaches aimed to understand if patterns found in long kept laboratory populations (pure inbreeding) are similar to those found in natural asexual populations as well as understanding how populations parameters vary between deferentially reproducing taxa.

To account for ploidy in the triploid asexuals, coverage ranges per genome copy were selected for all further analysis based on coverage distribution of the reads mapped against any of the reference genomes (supplementary figures 1-3). For (A) MAL data a coverage range of 15X per genome copy was selected (10X-55X for asexual triploid lines and 10-40 for sexual diploid lines). For (B) natural populations a coverage range of 23X per genome copy was selected (10X-80X for asexual triploid populations and 10-56 for sexual diploid populations). Pileup files were created individually for each genome data set of MALs and natural populations, respectively, using samtools mpileup (Danecek et al., 2021). A sync file was obtained using mpileup2sync.jar from Popoolation2 (version 1.201) (Kofler, Pandey, & Schlotterer, 2011) from all data sets together to maintain only all common positions, which served as input for the estimation of the fixation index *F*_*ST*_ on non-overlapping 1kb windows using Fst-sliding.pl. An approximate size of 3000 individuals per population was used.

*F*_*ST*_ was also estimated for 5 MAL of population size N=100 of *C. elegans* (Konrad, Brady, Bergthorsson, & Katju, 2019) from the Bioproject PRJNA448413 ([Dataset]Konrad, 2018) to compare our hermaphroditic sexual population to the model organism.

Individual pileup files were used as input for Variance-sliding.pl from Popoolation (Kofler, Orozco-terWengel, et al., 2011), using the options –measure theta and –measure pi respectively. The effective population size was then obtained as *N*_*e*_ = *θ/*2*nμ* for the sexual populations and asexual populations, where *μ* is the mutation rate estimated as described above and n corresponds to the ploidy of the studied system (n=2 for sexuals and n=3 for asexuals).

Theta of homologous genome copies (diploid part of the triploid genome) was estimated using Tetmer (version 2.2.1) (Becher et al., 2020). A k-mer spectrum of the Illumina reads per population was obtained using the K-mer analysis toolkit (KAT version 2.4.2) (Mapleson, Garcia Accinelli, Kettleborough, Wright, & Clavijo, 2016) (K=27) and provided to Tetmer as input. The manual fitting mode was used for both reproductive modes, for asexuals the triploid allopolyploid model (AAB) was selected, the diploid (AA) model was selected for sexual populations.

Plots for visualizing *θ*_*w*_, *π, F*_*ST*_ and effective population size results were obtained using the R package ggplot2 (Wickham, 2016). Statistical analyses to test for significance between the results obtained from each of the populations were done using Welch’s t-test. Welch’s ttest is used since sample sizes are unequal (more asexual populations than sexual populations) to compare the mean values of *F*_*s*_*t, π, θ*_*w*_ for the different reproductive modes.

#### 2.3.2 Phylogenetic network construction

A phylogenetic network was constructed for the parthenogenetic populations to test whether they can be defined as phylogenetically distinct species due to their genetic divergence. A phylogenetic network was also constructed for the sexual species as a reference. Orthologues present in all asexual and in all sexual populations were used for this analysis, each reproductive mode was assessed separately. Coordinates of the orthologue genes were obtained for each reference genome using Benchmarking Universal Single-Copy Orthologs (BUSCO) (Simão, Waterhouse, Ioannidis, Kriventseva, & Zdobnov, 2015). The coordinates of the genes on the reference genome extract single gene bam files, from alignments produced as previously described, for each population using samtools. To obtain a consensus sequence for each gene, bcftools (version 1.13) mpileup and bcftools call were used for variant calling (Danecek et al., 2021). These served as input for GATK (FastaAlternateReference- Make)(version 4.2.3.0))(Auwera & O’Connor, 2020) to obtain a consensus sequence per population for each gene, a multiple sequence alignment was obtained using MAFFT (version 7.47.1) (Katoh, 2002). A gene network was obtained for the consensus gene sequences of orthologues present in all the populations per reproductive mode using the median network algorithm from Splitstree4 (Huson, 1998).

## 3 Results

### 3.1 Low spontaneous mutation rate in parthenogens

We conducted a mutation accumulation line (MAL) experiment with an asexual (triploid) and a sexual population (diploid) (Figure 3). MALs were maintained for up to 40 generations, with many lines being lost over the course of the experiment. Starting with 100 and 71 lines respectively, 30 (30%) sexual lines and 15 (21%) parthenogenetic lines survived. Of these, six lines per reproduction mode were randomly chosen for whole genome sequencing and mutation calling. High-confidence mutations were called from 31,735 and 25,812 common positions in asexual and sexual lines, respectively. For the asexual lines, 72375 candidate mutations were called, whereas 7852 candidates were found for the sexual lines. After filtering for coverage ranges, read support, unique mutations per line, miss-mapped regions, and manual curation, 11 DNMs (De novo mutations) with high support remained for the asexual lines and 10 DNMs for the sexual ones (supplementary table 3).

Per reproductive mode five MAL showed at least one DNM. For the asexual lines an average of 7.5 *×* 10^10^ callable sites were found and 5.6 *×* 10^10^ for the sexual lines (supplementary table 3). This resulted in a total mutation rate of 5.828 *×* 10^*−*10^ (Bayesian HCI 1.2 *×* 10^*−*17^, 5.1 *×* 10^*−*9^) for asexual Panagrolaimus and a respective mutation rate of 8.923 *×* 10^*−*10^ (Bayesian HCI 2.9 *×* 10^*−*17^, 6.9 *×* 10^*−*9^) for the sexual Panagrolaimus. The mean rate for asexuals is lower than that found for sexuals, however, due to wide credibility intervals, mutation rates between asexual and sexual nematodes do not differ significantly. The majority of DNMs were transitions, with a transition/transversion ratio of 4.5 (9:2) in the asexual lines and 1.5 (6:4) in the sexual lines (supplementary table 5).

### 3.2 Different population genetic patterns between asexual and sexual lines under inbreeding

Standing nucleotide diversity, i.e. *π* of the respective parental lines, is lower in the sexual srefpool than in the asexual arefpool (Figure 4).

**Figure 4:**
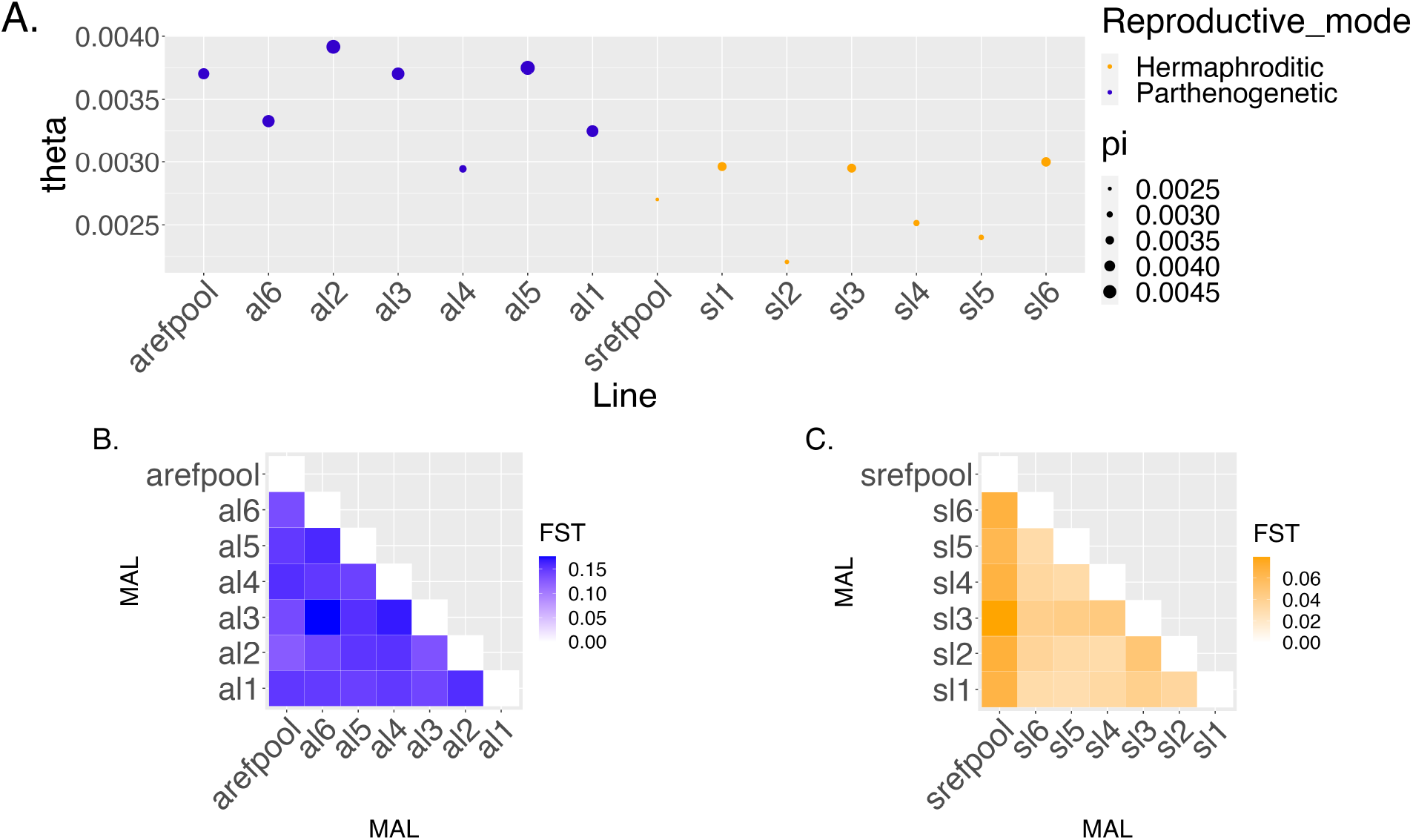
Nucleotide diversity (A) and population differentiation of (B) asexual and (C) sexual mutation accumulation lines. On average, *θ*_*w*_ and nucleotide diversity *π* are higher in asexual lines. A pattern of differentiation can only be seen among sexual lines. Note that the range of Fst valued differs between asexuals and sexuals.

Estimating population genetic parameters for MALs of both reproduction modes revealed significantly lower *θ*_*w*_ (*p <* 0.0005) in sexual lines (range 0.0022-0.0029 - lowest and highest *θ*_*w*_ values found in the different sexual lines)(supplementary table 6) when compared to the asexual lines (range 0.0029-0.0037 - lowest and highest *θ*_*w*_ values found in the different asexual lines). Sexual lines are also significantly lower in nucleotide diversity (range 0.0024- 0.0036)(*p*value = 0.001315) with the lowest value found in the sexual parental population srefpool, when compared to asexual lines. Nucleotide diversity of asexual lines ranges between 0.0032 and 0.0046, with the lowest *π* found in one of the asexual MAL (al4), which is also the asexual line with the lowest *θ*_*w*_.

Comparisons of population differentiation among parental and descendant lines per reproduction mode revealed distinct differences. The sexual lines showed a significantly higher *F*_*ST*_ between parental and descendant lines when compared to the *F*_*ST*_ among descendant lines (*p*-value== 6e*−*7)(supplementary table 8). For the asexual lines, this comparison was not significant (supplementary table 7). In general, genome-wide population differentiation was found to be more pronounced among asexual lines (including parental arefpool - PS1159 parental) with *F*_*ST*_ ranging between 0.1312 and 0.1755 in comparison to the *F*_*ST*_ range in sexual lines (0.0304-0.0796). Population differentiation was significantly higher in asexual lines than in sexual lines (*p*-value =7.5 *×* 10^24^). *F*_*ST*_ values found on the sexual descendant lines were similar to those estimated for *C. elegans* descendant lines (population size 100)(supplementary table 9) from a MAL experiment performed by (Konrad et al., 2019).

### 3.3 High genetic diversity but low differentiation in natural populations of parthenogens

Nucleotide diversity *π* for asexual populations ranged from 0.0402963 (Raleigh, USA; apop2) to 0.206227 (Beijing, China; apop3). For sexual populations *π* ranged from 0.00241247 (Guangxi, China; spop4) to 0.0545159 (Cologne, Germany; spop3). Nucleotide diversity was on average higher for asexual populations (*p*-value= 0.0145)(supplementary table 10) (Figure 5) .

**Figure 5:**
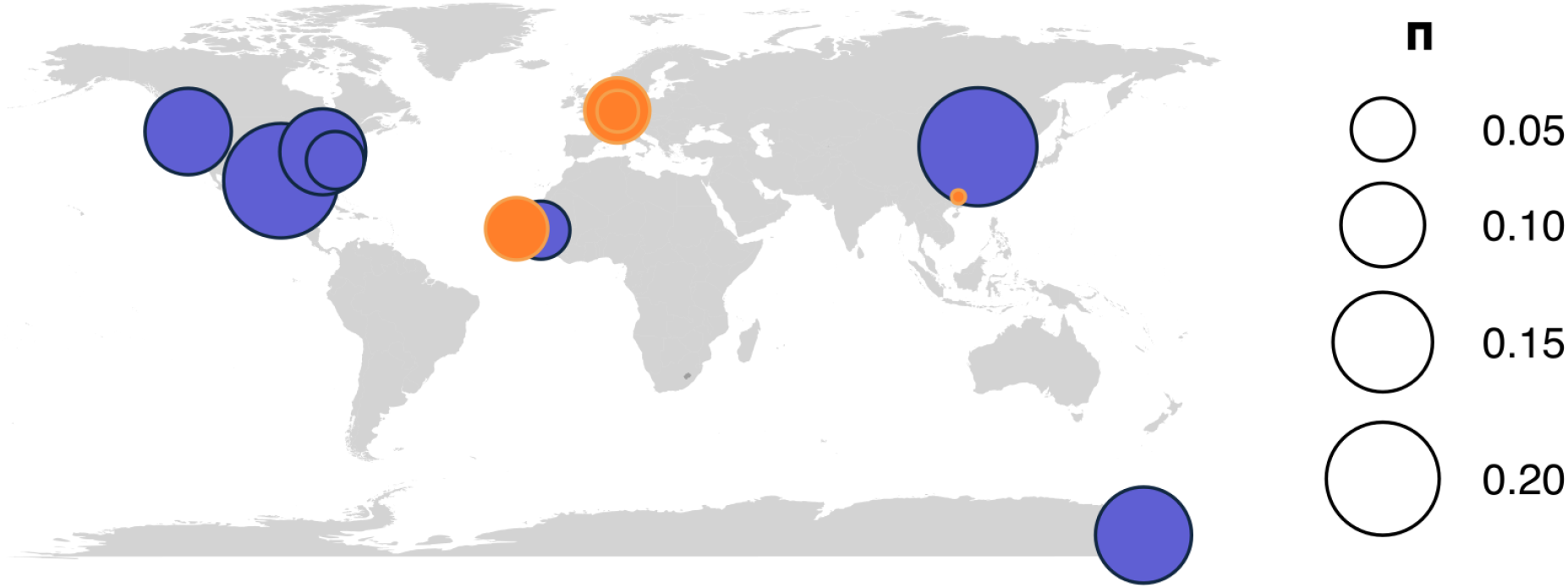
Nucleotide diversity *π* on natural populations of nematodes isolated from distant geographical areas. Nucleotide diversity is higher on natural asexual populations when compared to natural sexual populations

Genome-wide estimator *θ*_*w*_ for asexual populations ranged from 0.0306732 (Diourbel, Senegal; apop1) to 0.192175 (Beijing, China; apop3). For sexual populations *θ*_*w*_ ranged from 0.0027009 (Guangxi, China; spop4) to 0.0452624 (Cologne, Germany; spop3). Asexual populations showed a significantly higher *θ*_*w*_ when compared to sexual populations (*p*-value = 0.0242).

To test whether the higher diversity found on asexuals could be caused by the third genome copy, estimator *θ*_*w*_ was obtained for homologous genome copies (asexual triploid *Panagrolaimus* have a hybrid origin). For triploid asexual populations, the mean *θ*_*w*_ obtained for homologous copies (2 genome gopies) was generally lower than genome wide *θ*_*w*_ estimations (3 genome copies), but not significantly different (*p*value = 0.0747)(supplementary table 11).

In natural populations, population differentiation *F*_*ST*_ between asexual Diourbel population, Senegal (apop1) and asexual Corvallis population, USA (apop7) was highest with an *F*_*ST*_ of 0.468 for asexual populations (supplementary table 13). For sexual populations, the highest level of population differentiation was found between the Santiago Island population, Cape Verde (spop1) and the Cologne population, Germany (spop3) with an *F*_*ST*_ of 0.851948 (supplementary table 12). Population differentiation was more pronounced among sexual populations as compared to among asexual populations (*p*value=0.00002471).

### 3.4 Asexual *Panagrolaimus* populations are genetically distinct to each other

To analyse divergence between the monophyletic asexual *Panagrolaimus* populations under the phylogenetic species concept we used a network approach in Splits Tree (Huson, 1998). In the asexual reference genome (*Panagrolaimus sp*. PS1159) we idendtified 2173 universal single-copy orthologues. For each population, reads corresponding to the coordinates of the orthologues were extracted in order to generate consensus sequences for genes with sufficient read support: 375 orthologues were shared on the asexual populations, these genes were aligned and used on the gene network. We conducted a corresponding analysis for the sexual species as proof of principle. For the sexual reference (*Panagrolaimus sp*. bornheim) genome 1983 orthologues were found and reads corresponding to these genes’ coordinates were extracted; a total of 213 orthologues were used for the gene network. The Median network calculated in Splits Tree showed a total of 23 splits between the asexual populations, while there were 3 splits between the sexual species, i.e. the minimum possible number of splits (Figure 6).

**Figure 6:**
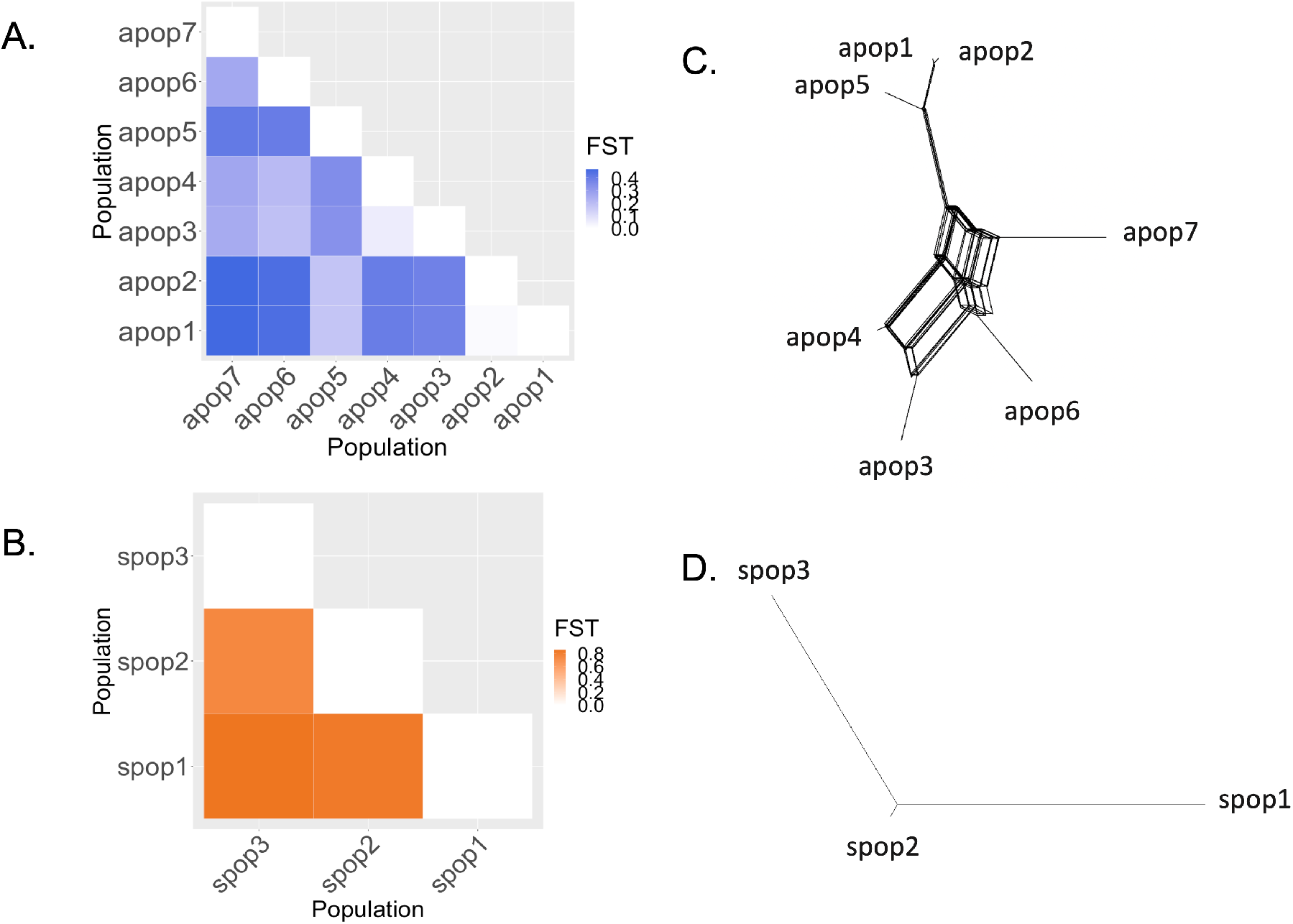
Population differentiation of (A) natural asexual populations and (B) natural sexual populations. Orthologue gene network of (C) asexual and (D) sexual populations of *Panagrolaimus* nematodes. Sexual populations show higher differentiation than asexual ones. The gene network shows that asexual nematodes analyzed are genetically distinct from each other.

## 4 Discussion

In this study we aimed to analyze divergence patterns in the evolutionary context of asexual animals using parthenogenetic *Panagrolaimus* nematodes as a system, and sexual, in this case fully selfing hermaphroditic *Propanagrolaimus*, as a comparator. By running long-term MAL we conducted a classical experiment to measure mutation rates under nearly neutral evolution(Halligan & Keightley, 2009). We also estimated standard population genetic parameters from natural populations of widely-spread geographic origin (Figure 5)(gonochoristic sexual *Panagrolaimus* and parthenogenetic asexual *Panagrolaimus*), and used phylogenetic networks (Figure 6) to narrow the complexity of population genomic patterns under parthenogenesis.

Genome sequencing has seen another drastic advancement in the last years with longread methods becoming available for single individuals with sufficient amounts of DNA (e.g. vertebrates), thus potentially allowing for higher resolution in population genomic assays (De Coster, Weissensteiner, & Sedlazeck, 2021). In our study we still depend on shortread sequencing data of pools of individuals, as single individual long-read sequencing from tiny organisms remains challenging. Consequently, our study remains reference-based with limited resolution on aspects such as structural genomic variation (Adewale, 2020). At the same time, short-read (Illumina) sequencing still outperforms long-read methods in terms of the cost-coverage-ratio when aiming for deep sequencing studies, which is inevitable for the here conducted MAL experiment for the identification of *de novo* mutations.

### 4.1 Lower spontaneous mutations rate in parthenogens could aid to diminish the effect of Muller’s ratchet

Parthenogenesis is assumed to have low evolutionary potential due to the accumulation of deleterious mutations (Muller’s Ratchet) (Muller, 1932), lack of recombination and low genetic diversity (Simon et al. 2003). In *Panagrolaimus*, the parthenogenetic populations analysed here appear to be monophyletic, originating 1.3–8.5 Mya ago through hybridization involving triploidization (Schiffer et al., 2019). Whereas sexual populations and species in our analysis undergo normal meiosis, eggs of the nematode *Panagrolaimus sp*. PS1159 (arefpool and apop2) develop without fertilization and the offspring is exclusively female. Asexual meiosis (presence of polar body) occurs without recombination [pers. comment, Caroline Blanc and Marie Delattre, Lyon] and as a result, offspring are clones of their mother. The majority of non-neutral mutations that occur are deleterious (Sloan & Panjeti, 2009). To explain long-term persistence of asexuals it can be assumed that they have lower mutation rates, thus the process of mutation accumulation is slowed down. Although the estimated mutation rates did not differ significantly between sexual and asexual *Panagrolaimus* populations, genetic diversity in the MAL was found to be much higher in asexual *Panagrolaimus* than in sexual *Propanagrolaimus*.

If mutation rates are low and recombination is absent, clonal interference can be reduced when beneficial mutations in a population arise. With higher mutation rates, if many beneficial mutations are carried in different clones, fixation time of these mutations is increased (De Visser & Rozen, 2005). In our study, mutation rates found for both reproductive modes are one order of magnitude smaller than that reported for the distantly-related nematode *C. elegans* (Denver et al., 2009) and for the arthropod model *Drosophila melanogaster* (Keightley, Ness, Halligan, & Haddrill, 2014). This not only shows that findings in model organisms cannot be generalized across phyla, but also indicates divergent evolutionary mechanisms in both panagrolaimid nematodes analysed here. To better understand these, it will be necessary to estimate mutation rates with additional MALs, run for shorter accumulation periods to reduce variation of the data and thus allowing for better statistical validation (Oppold & Pfenninger, 2017).

### 4.2 Different population genetic patterns between asexual and sexual lines under inbreeding

Asexual reproduction may affect population differentiation patterns differently than sexual reproduction does (Balloux, Lehmann, & de Meeûs, 2003; Prugnolle & De Meeûs, 2008). Theoretical examples of clonal (=asexual) diploid parasitic species, show that populations are expected to be less differentiated in comparison their parents (i.e. preceeding generations) than to other populations (Prugnolle & De Meeûs, 2008). In our study the triploid asexual MAL did not produce a distinct pattern of differentiation between MA lines and the parental state or among MA lines (Figure 4). For sexual reproduction, previously described highly inbred laboratory selfing nematodes, such as *C. briggsae* and *C. elegans*, show low heterozygosity and a higher differentation between populations and the parental state than among populations, which is a result of alternative alleles being fixed separately in each lineage (Barri`ere et al., 2009; Teotónio, Estes, Phillips, & Baer, 2017). Our results for the sexual (hermaphroditic) nematodes align with said “ sexual “pattern: reduced heterozygosity.

Parthenogenetic *Panagrolaimus* nematodes analysed in this study have been found to be triploid by (Schiffer et al., 2019) and other asexual representatives of the genus are also triploid (Shatilovich et al., 2023). The lower rate of differentiation among sexual lines from the MAL experiment in comparison to asexual lines could be explained by (i) lower standing genetic variation in the hermaphroditic parent at the beginning of the experiment and (ii) increased heterozygosity for asexuals arising from the third genome copy. In asexual lines “inbreeding” does not change the genetic structure of a population (Bengtsson, 2003). The pattern of population differentiation we found for asexual MA lines could show the differentiation pattern of natural populations: maintained heterozygosity within offspring due to the lack of allele segregation (Stoeckel & Masson, 2014). It has been proposed that under asexuality, heterozygosity can even increase since alleles of a same gene could independently accumulate mutations over generations, the so-called Meselson effect. This has recently been observed in obligate asexual oribatid mites (Brandt et al., 2021), but could not be tested in the nematodes with our current data.

### 4.3 High genetic diversity but low differentiation in parthenogenetic populations

Studies on populations of the sexual parasitic nematode *Baylisascaris schroederi* isolated from different mountain ranges, showed very low and non-significant *F*_*ST*_ (range: 0.01911 to 0.02875) between three populations (Zhou et al., 2013). Natural populations of sexual *C. brenneri* also show low values of differentiation between populations, in this case, from geographically distant regions eastern India and French Guiana (0.092)(Dey, Chan, Thomas, & Cutter, 2013). Our results reveal contrasting patterns of on average higher differentiation among natural sexual populations. In our study, both sexual and asexual populations have two-fold higher *F*_*ST*_ values than those reported for *Caenorhabditis* nematodes and *Baylisascaris schroederi*. These higher *F*_*ST*_ values could be an initial indication for local adaptation (Hirase et al., 2021) to the distinct regions where the populations were isolated from.

Population genetic parameters obtained for asexual natural populations showed a mean nucleotide diversity higher than that found for sexual populations. This pattern contradicts the expectation that the lack of recombination is assumed to result in reduced genetic diversity in parthenogenetic taxa compared to sexually reproducing counterparts (Charlesworth & Willis, 2009). This conflictual pattern of increased nucleotide diversity has also been found for the amazon molly, an asexual fish of hybrid origin (Jaron et al., 2021), and in the asexual bark lice (*Echmepteryx hageni*) (Shreve, Mockford, & Johnson, 2011) when compared to respective sexual counterparts.

*θ*_*w*_ was also found to be higher in the asexual populations genome-wide estimations. However, when only homologous genome copies were compared (accounted for by allopolyploid model using Tetmer), *θ*_*w*_ estimations were generally lower, revealing the third genome copy as a major source for genetic diversity due to the hybrid origin. Increased genetic diversity in asexuals could also be explained by a larger effective population size *N*_*e*_ (Soulè, 1976). Under the neutral theory, genetic diversity is expected to increase with *N*_*e*_ and this has been seen in taxa such as chordates, annelids, arthropods and molluscs among others (Buffalo, 2021; Leffler et al., 2012). Larger, more stable populations are then expected to maintain greater levels of neutral genetic diversity than populations with lower effective population sizes *N*_*e*_ (Leffler et al., 2012).It is predicted that large population sizes in asexual taxa aid the avoidance of extinction since the effectiveness of natural selection is increased and the lack of recombination can be compensated in such lineages (Ross, Hardy, Okusu, & Normark, 2013). Consequently, studies have found larger populations for asexual oribatid mites than in sexual ones in temperate and boreal forests (Brandt et al., 2017).

Heterozygosity is heavily affected by effective population size. This could differ between asexual and sexual populations because of differences in actual population size, reproduction mode and the different impact of purifying selection. For example, an increased mutational load in asexuals could reduce their effective population size due to purifying selection, but this effect could be compensated by larger population sizes.

Asexual reproduction accompanied by high genetic diversity has shown to allow for rapid adaptive responses on parasitic nematodes of *Meloidogyne* species to their hosts (Castagnone- Sereno, 2006). Nucleotide diversity of natural *C. elegans* populations from Hawaii has been found to be lower than that found here for asexual and sexual populations, these Hawaiian isolates are known to harbor a higher genetic diversity than all other known *C. elegans* populations. However, the average genome-wide diversity found for these *C. elegans* populations (*π* 0.00109) is very similar to what was obtained for JU765 (spop4 - 0.00129). Both *C. elegans* and our *Propanagrolaimus* populations are diploid and selfing hermaphrodites (Crombie et al., 2019). The variable proportion of nucleotide diversity found on different genomic regions, is consistent with the assumed pattern across the genomic landscape. Some regions have very low diversity and could correspond to coding regions, whereas introns are expected to be more diverse (Tatarinova et al., 2016). Follow-up studies, including the annotation of reference genomes, will allow for more precise estimations of *π* across the genomic landscape.

### 4.4 Asexual *Panagrolaimus* populations are genetically distinct to each other

Split networks have been successfully applied to compare distantly related taxa (yeast, mammals, *Drosophila*, and *C. briggsae*)(Huson & Bryant, 2006), as well as populations of hyperdiverse nematodes (Dey et al., 2013). Our split network analysis based on 375 orthologues in the asexual populations and 213 orthologues in the sexual species, appears to indicate that the former are as genetically distinct as the latter. Notably, the network for the sexual species is tree-like, while it shows more splits for the parthenogens. This is expected in a triploid system, where homeologs are not resolved (or phased in the genome assemblies) and thus recapitulate a pattern usually seen in recombining populations.

The *Panagrolaimus* genus is characterised by very little morphological variation, usually only observable under the electron microscope to the (taxonomic) expert eye. Thus, morphological species descriptions are limited and isolates are referred to as strains. Parthenogenetic taxa add an extra level of complexity as the classical biological species concept, which defines species as groups of (potentially) interbreeding populations which are reproductively isolated from other similar groups (Mayr, 1999), is not applicable. However, distinct species have been found for asexual organisms such as bdelloid rotifers, oribatid mites and oligochaete worms by using genomic data (Birky, Adams, Gemmel, & Perry, 2010), i.e. applying a phylogenetic or phylogenomic species concept.

We had initially decided to treat the different nematode isolates/strains as separate populations according to their geographical origin and reproductive mode. While analyzing population differentiation (Fst), it became clear that the populations being analyzed showed higher Fst values than other nematode populations previously studied. Hence, we tested how genetically distinct the different populations were to obtain insights on whether they could be genetically distinct species despite the low morphological variation and lack of interbreeding, using single-copy orthologues.

### 4.5 Theoretical expectations about mutation rate evolution need to be adapted to diverse and complex systems

Mutations are the ground source of genetic variation, even if most non-synonymous mutations that occur are deleterious. While both mutation and recombination rates are variable along the genome, in obligate asexual taxa recombination can be completely absent, possibly leading to the accumulation of mildly deleterious mutations. We have found indications that natural populations of asexual *Panagrolaimus* show an elevated level of heterozygosity, potentially due to their third genome copy and the lack of allele segregation. The mutation rate in these organisms appears to be low, thus maybe delaying the effect of Muller’s ratchet. The differentiation found within sets of natural populations, as well as the genomic differentiation between populations seems to show the potential for evolution in parthenogenetic animals. Asexual reproduction does not occur in the same way across the tree of life. In some taxa like asexual angiosperms, asexual reproduction occurs with meiosis still present and where gene conversion can occur. In other taxa, such as *Panagrolaimus* nematodes, asexual meiosis happens without recombination, whereas in other nematode species (Castagnone-Sereno & Danchin, 2014) and stick insects (Schwander, Henry, & Crespi, 2011), mitotic parthenogenesis without meiosis can take place.

### 4.6 Conclusions

Asexual reproduction adds another level of complexity to our understanding of mutational evolution, selection, and adaptation. Many expectations for genome evolutionary processes, as e.g. the reproductive mode violates the classical population assumption of random mating (Fisher, 1923)(Wright, 1931). In our study it became obvious that studying “parthenogenomic” complexity is currently limited on two very different levels. The availability of genomic resources is limited (phased reference genomes from tiny invertebrate organisms are not available for enough representatives of the Phyla). Polyploid genomes, as those in the asexual populations studied here, are technically challenging as they are likely to involve divergent evolutionary trajectories between homeologs, which carry different patterns of variation (Hörandl et al., 2020). To better infer measures such as *π* phased reference genomes should be used and with the current advances in long-read sequencing methods these should be available in the future (The Darwin Tree of Life Project Consortium et al., 2022) even for tiny invertebrate taxa, as nematodes or rotifers. In this study we aimed to account for potential biases by adjusting coverage ranges to ensure high enough representation of the respective genotypes in the short-read data and thus allow for each position on each genome copy to be called equally likely. However, to study such genotypes on a single individual level in polyploid systems it is urgently necessary to develop more sensitive software tools. As parthenogentic taxa are not paying the “cost of sex”, they could have an advantage in extreme, and challenging environments (geographical parthenogenesis)(Tilquin & Kokko, 2016). This pattern has been observed in plants, flatworms (Lorch, Zeuss, Brandl, & Brändle, 2016) and ostracods (Symonová et al., 2018). Based on our findings that genomic diversity is high in asexual populations, we would like to extend our research to further analyse a large set of populations isolated from distinct extreme environments to test the hypothesis of geographical parthenogenesis.

## 5 Acknowledgments

The authors gratefully acknowledge the help of Michael Kroiher and Einhard Schierenberg in conceiving the original MAL experiment. The authors are thankful to Tanja Schwander for comments on analyses methods, and to Caroline Blanc and Marie Delattre for a cytological analysis in two *Panagrolaimus* nematodes. This project was supported by, initially, a grant from the Volkswagen Foundation to P. Schiffer. The project was then supported through a DFG ENP Projekt [grant number 434028868] to P. Schiffer, and the DFG funded project B08 in the CRC1211 [grant number 268236062] to P. Schiffer and A-M Waldvogel, in which L. Villegas is employed. We furthermore thank the Regional Computing Center of the University of Cologne (RRZK) for providing computing time on the DFG-funded [Funding number INST 216/512/1FUGG] High Performance Computing (HPC) system CHEOPS as well as support.

## 6 Data Accessibility and Benefit-Sharing

Genome assemblies and sequencing data are deposited under Bioproject PRJNA374706 and are available through Wormbase.

The authors confirm contribution to the paper as follows: study conception and design: P. Schiffer, T. Wiehe; data collection: Y. Author; analysis and interpretation of results: L. Villegas, L. Ferreti, A. Waldvogel, P. Schiffer; draft manuscript preparation: L. Villegas, L.Ferreti, T. Wiehe, A. Waldvogel and P. Schiffer. All authors reviewed the results and approved the final version of the manuscript.

## 7 Tables and figures

